# Detailed curation of biological samples and experimental designs for genomics using LLM-supported agentic workflows

**DOI:** 10.64898/2026.07.30.741874

**Authors:** Paul Pavlidis, B. Ogan Mancarci, Amanda Maximo, Carlton Yan, Rachel A. Schwartz

**Affiliations:** Department of Psychiatry and Michael Smith Laboratories, University of British Columbia, 2185 East Mall, Vancouver BC V6T1Z4 Canada

## Abstract

We describe an automated software tool to accomplish data curation tasks previously performed by humans for the Gemma genomics data re-analysis resource. Gemma is a hand-curated database of reprocessed transcriptomic studies, currently covering over 23,000 human, mouse and rat data sets largely drawn from the Gene Expression Omnibus (GEO). We developed a pipeline that uses both traditional (mechanical) and large-language models to produce detailed ontology-anchored, sample- and experiment-level annotations in accordance with our established curation guidelines. In this report, we describe benchmarking the pipeline and investigations aimed at evaluating readiness of the v1.1 Gemma curation agent for production use. Overall, performance is near that of human curators, at approximately 1/20th the cost and at least 100 times the speed. We also present preliminary exploration of triage methods for identifying agent curations that are more likely to contain errors, and thus can be forwarded for human review. We discuss the potential place of such curation approaches in bioinformatics ecosystems. Besides the software, our deliverables include the benchmark set of 500 studies and an evaluation framework that can be used to further develop the pipeline or compare to other approaches.

## Introduction

Manual data curation has historically been a major task in biomedical knowledge engineering, and one that is increasingly tantalizing to automate using artificial intelligence (AI) and large language models (LLMs) in particular. Here we provide a preliminary report on our recent work to develop and implement approaches to automate detailed experimental metadata for genomics research.

The context of our work is the Gemma database (<u>gemma.msl.ubc.ca</u>). In operation since 2005, Gemma has accumulated over 23,000 transcriptomics studies, mostly sourced from GEO (Clough et al. 2023). We have hand-curated them to describe the experimental design and other features of the data at the sample level (**Figure 1**). Where possible terms from ontologies are used, and the process is guided by an extensive and detailed curation handbook. There are some automated steps and many software assists to improve curator efficiency and accuracy, but the process remains laborious (Lim et al. 2021). With the recent versions of our pipeline and a team of 2-3 personnel, we were able to curate up to 2,000 or 3,000 studies per year. However, while Gemma has high-quality annotations overall, there are inconsistencies, errors and gaps. This motivated us to explore how we can improve past annotations as well as for new data.

**Figure 1:**
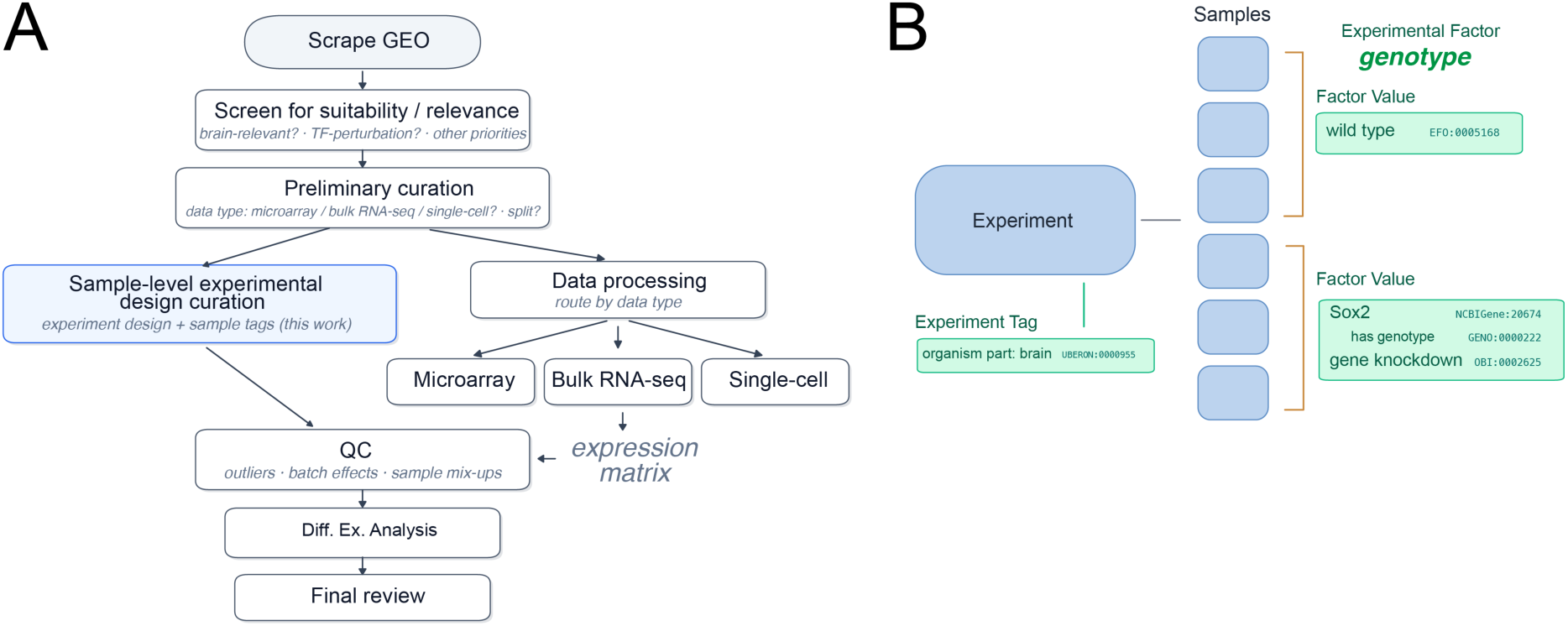
The curation task and the data model. **A**: The curation workflow for a gene-expression series, from scraping GEO, screening for suitability, and preliminary curation, through sample-level experimental-design curation (blue; the focus of this work), to data processing by assay type, quality control, differential-expression analysis, and final review. **B**: The basics of the curation data model; green indicates ontology-grounding. An experiment owns its samples and carries whole-experiment tags (e.g. organism part: brain) and experimental factors (e.g. genotype). Each factor value is described by ontology term statements and applies to a specific set of samples.

Automating data curation has long been an active area of research, with varying degrees of success. In work early in Gemma’s lifetime, we developed methods to do one specific task: to tag each experiment with ontology terms to capture basic information such as the tissue studied, mouse strains, disease and cell lines, to complement the sample-level annotation (French et al. 2009). Benchmarking against human curators, we were able to achieve a recall of about 0.5 but a precision of only 0.15-0.25. While helpful, it was not good enough to use without human intervention to fill in gaps and remove excess tags. We soon returned to manual methods, as the automated methods were also too primitive to account for the subtleties of our curation guidelines, which also evolved.

Until recently, the more ambitious task of automating the full detailed curation of sample characteristics was out of reach. This is because the tasks required human judgement, both to find obscure pieces of information buried in a supplementary file or methods section, or cryptically encoded in sample names, but also to align the curation with our complex and sometimes idiosyncratic guidelines. It typically takes months to train curators, and proficiency increases further over a longer period. Even our most experienced curators make errors, and while we conduct cross-checks, inconsistencies and gaps are present. Even in the absence of errors, and even constrained by detailed rules, curation is a sufficiently flexible task that there is room for valid annotations to be expressed in different ways. Our own experience is highly consistent with the literature (**Supplementary Figure 1**): inter-curator agreement is around at most 90%, and depending on the granularity and degree of judgement required and flexibility in the task, it can be substantially lower (the tasks we address in this paper range from ∼60%-90% curator agreement).

The design of Gemma’s curation framework was heavily influenced by limitations in resources, and was designed to not be too difficult, mentally draining, nor time consuming to do correctly. This means we made compromises that we would not have made if the process was fully automated from the start. While many people enjoy data curation, there is a consensus among those who have worked on Gemma that it can be repetitive and is not always intellectually engaging. They have always welcomed, promoted, and developed ways to streamline and automate their work. To this end we have long worked to improve our software tools, including adding a step in our intake process to fill in some ontology terms automatically. These and other changes probably led to a doubling of throughput in aggregate, but human effort remains the bottleneck.

In the last few years, we have seen LLMs emerging as supporting tools for data curation. We recently developed an approach to identify mouse strains and cell lines in genomics studies that were used as the source of the samples, and retrieve the correct ontology term (Rogic et al. 2026). We achieved state-of-the art (but well short of perfect) performance and also identified errors in Gemma as well as in the underlying GEO records. Other previous work with LLMs showed promising results for a range of data annotation and information extraction tasks (Gue et al. 2024; Mahmoudi et al. 2024; Chen et al. 2025; Jensen et al. 2025; Kainer 2025; Poretsky et al. 2025; Riquelme-García et al. 2025; Turner et al. 2025).

Recently (Mittal et al. 2026) used an LLM to identify cell types, tissues and other features in GEO studies as part of a system called MetaMuse, which we previously evaluated (Rogic et al. 2026). A few other groups have also built agentic pipelines for GEO or SRA metadata. (Mondal et al. 2025) use a six-stage multi-agent system to curate single-cell metadata fields, and (Hak et al. 2026) combined rule-based extraction with a fine-tuned LLM for SRA cancer metadata. These systems target a sub-part of our curation task. The need for Gemma is arguably more complex, as it requires understanding the structure of the experiment and finding information in disparate sources, and then documenting the salient details according to specific set of criteria and ontologies.

The other reports and our own findings on strain and cell type resolution were encouraging, but the errors led us to anticipate, even as recently as a few months ago, that humans would probably have to review much AI curation. Two things have shifted our thinking. First, it is apparent that the latest LLMs (after late 2025) have capabilities that make attempting much more ambitious tasks seem worthwhile, especially for writing code but also for natural language processing. Second, we don’t think having humans review machine-generated curation is desirable. If every item has to be reviewed by a curator, even briefly, the time savings will still be real, but the enjoyable aspects of data curation will be sapped, turning the human from one who is empowered by a tool into a “reverse centaur” whose job is instead defined by the AI (Doctorow 2026). A goal of the work we describe here is to reduce the need for human review to an absolute minimum; anything short of that would not be enough of an improvement to get excited about. The system we aim for would handle most of the work without intervention, and forward only difficult cases for human input.

Here we show that while some challenges remain, it is now possible to heavily support, and even replace, human curators with AI for a large fraction of the curation tasks needed for Gemma. Quantitatively and qualitatively, performance is very close to human. There are errors, but we consider them likely tolerable as a trade-off for efficiency gains and side-benefits such as provenance and supporting evidence for every annotation, especially if we can effectively triage human review. We still expect the transition to near full-automation to have some bumps, and we discuss reasons for both optimism and caution for the future of biomedical data curation.

## Methods

### Overview of GEO and Gemma

We assume the reader is somewhat familiar with GEO and the meta-data provided in a Series (Identified by a GSE ID such as GSE123) and its associated Samples (GSM IDs). Gemma supports transcriptomics data from microarray, RNA-seq, and single-cell platforms. We provide an overview of Gemma’s curation practices in **Figure 1** and summarize some salient information on both GEO and Gemma in this section.

GEO records conform to a schema which suggests many fields that can be provided by data submitters, as free text. The schema has evolved over the years, but the fields remain mostly free-form and not mapped to ontology terms. To our knowledge, there is no curation conducted by GEO and the submitted annotations are generally presented as-is, which is why we developed Gemma.

Gemma imports the GEO “Sample characteristics” as “Biomaterial characteristics”, and they form a primary starting point for curation along with the rest of the GEO metadata (Overall design, Extraction Protocol, etc., when present) and the source publication (if available). The main curation task is to more formally define the properties of the samples which pertain to biological signal. This has no limit of variety, but Gemma broadly apportions these features into categories such as “genotype”, “cell line”, “organism part”, “disease”, “strain”, “cell type”, “treatment”, “developmental stage” and “biological sex”. The information might be in the GEO sample characteristics, other meta data such as the overall design, or the publication. A single GEO sample characteristic can describe multiple features (e.g., “male mouse treated with drug”) that need to be parsed, and can be very cryptic. In Gemma, we treat features that vary across samples as (potential) *Experimental Factors* that have levels defined either as continuous (such as mass) or (more commonly) categorical *Factor Values* (**Figure 1B**).

Features that are constant across all samples are also annotated. In Gemma, for mostly historical reasons these are manually added as *Experiment Tags*, rather than to each sample (**Figure 1B**). However, our guidelines around what qualifies as a Tag has evolved over the years, leading to some inconsistency. Our current guidelines are to *not* annotate what the experiment is about or “relevant to”, but some such legacy curation is still present. Another relatively recent improvement to Gemma was to allow annotation to take the form of not just a category and an ontology term (“treatment [EFO:0000727] = bortezomib [CHEBI:52717]”), but more complex statements (subject–predicate–object), modeled on the Resource Description Framework (RDF) (Cyganiak et al. 2014). This allows Gemma to express (with some compromises) concepts such as “bortezomib [CHEBI:52717] delivered at dose [TGEMO:00166] 10mg/kg for 1 week” or “astrocyte [CL:0000127] located in [RO:0001025] cerebellum [UBERON:0002037]” (Figure 1B has another example). This has not been fully backported to older annotations as it often required human intervention to do so accurately (only 10% of annotations in Gemma have at least one predicate, though far from all need to exercise this pattern). We use terms from a set of 13 widely-used ontologies (see Supplementary Methods), and any gaps in ontology support are either left as free text or (more rarely) added as concepts to our in-house ontology (https://github.com/PavlidisLab/TGEMO).

As of the start of the work we describe here, we had over 23,500 transcriptome studies for mouse, human and rat, drawn from over 22,800 GSEs (a total of over 770,000 samples). While there are some non-GEO studies in Gemma, the main reason there are more studies than GSEs is because many series contain data from more than one species, or with two incompatible technologies, leading to a *split*, which in Gemma is indicated by a numeral after the GSE ID such as GSE1234.1 and GSE1234.2 (largely these splits are made automatically by Gemma but occasionally it is done manually). Some curation is done automatically, primarily assigning ontology terms for common textual descriptors associated with sample characteristics in GEO (biological sexes, some tissues, cell lines and mouse strains - a total of ∼1300 terms). Some additional details are provided in the relevant sections below.

### Overall approach

Our goal was to replicate our curation process and guidelines in an automated tool. We decided to do this, not because we think our curation guidelines are optimal, but because it gave us a fixed objective and let us leverage the curation we had already done. The entire study was conducted using Anthropic’s Claude Code (version 2) to develop the framework, design and conduct evaluations, and assemble the resulting data. We used Opus 4.7 or 4.8 as the coding model, at high effort (1M token context).

The first step was to export the relevant parts of our curation guidelines manual from our internal web site (161 pages, ∼104,000 words, 147 figures, 11 screen recordings). This, along with the Gemma code base and simple prompting was used to direct Claude to create a distilled version of the guidelines (which were somewhat chaotically organized and contained some old information and contradictions). We reviewed these guidelines manually and made minor adjustments because some rules were embedded in code or had not been fully documented in the first place. The richer statement-style annotation that replaced our older bag-of-terms approach was an example of such a gap. But we resisted the temptation to materially change the guidelines during development.

We used Claude Code to develop a system that conducts the same tasks as the curators and adheres to the same guidelines. This took the form of two main code repositories: *gemma-curation-agents* which is the implementation of the curation tasks and *gemma-ui*, which includes a new curation user interface (UI) that combines our existing manual editing capabilities with computationally-provided proposals side-by-side (**Supplementary Figure 2**). We designed a new UI because our existing one is showing its age, and trying to add such a major new feature would add to the challenge. The curation proposer is over 100,000 lines of Python, driven by 24 prompt and rule files totaling roughly 50,000 words. Bulk LLM calls are issued through the Anthropic Batch application programming interface (API) with prompt caching, which served the shared rules and system prompts from cache on about 60% of calls. Other models were accessed via a separate commercial provider (Together.ai).

A guiding principle for development was that as much as possible, we would rely on non-LLM methods. We loosely refer to these as “deterministic” or “mechanical”. This is related to the next principle, which is that everything needs provenance. For mechanical methods, that accounts to the script and the settings used to call it as well as inputs and outputs. For LLM calls, the model is always prompted to provide supporting quotes and/or justifications, which are carried forward to later stages as inputs but importantly contribute to verifiability. Another principle was to leverage the existing curation in Gemma. The main way this came into play (outside of evaluation) was to provide information on previous usage of ontology terms in the system. This was already used by curators to improve harmonization (by using terms already in use rather than new synonyms) and correctness (in cases of ambiguity, a matching term we previously used might be more plausible).

The development process was to rapidly iterate through two steps: first, write (or modify) code to do some aspect of the curation task and build up the pipeline, and second, try it on one or a few experiments and compare the results to the Gemma curation. At first the evaluation was very imprecise, in part due to the use of “eyeball” scrutiny at the earliest iterations, but it also emerged that comparing two sets of annotations is non-trivial and it took quite a few iterations to get right (and still is being improved). The coding agent made this process very rapid and allowed us to test dozens of iterations of the pipeline. At each iteration, the first author made a decision, often informed by input from Claude, as to how to improve. As we advanced and solidified the framework, we began more formal evaluation of larger numbers of experiments (see next sections). Claude was used to assist in data analysis and in ad hoc evaluation, such as helping sort through different types of errors and characterizing them.

Observations of the agent results on training data informed adjustments to the pipeline. To give an example from early in the process, the agent was mistaking a shorthand for Zeitgeber time (relative to the daily dark-light cycle) as a made-up genotype (“Z8”). Zeitgeber comes up in experiments occasionally, was explained in the GEO meta-data, and was already addressed in our guidelines, so we made adjustments to both prompts and mechanistic steps to better accommodate time points expressed in such ways. This proved effective and we saw little reason to think this wouldn’t generalize to future cases. Such corrections, on a small and large scale, were repeatedly implemented, with the coding agent always instructed to embed general principles, not an endless series of special cases. Despite this, the system is complex with numerous details expressed in code and prompts. We went through multiple rounds of code and prompt review and cleanup, but at this stage of development, the system likely has bugs and brittleness yet to be exposed, as well as easy ways to improve (in fact, the experiments we report are already becoming out of date as we move past v1.1).

### Training and evaluation data

We defined three sets of studies from Gemma: a validation set of 2,394 previously curated studies (chosen randomly with a bias to more recent studies) that were completely held out for later testing; 17,404 that were made available to provide context and which could be used for making indexes and embeddings, and another ∼2,200 studies available to the method development process. Most of the 2,200 we not used yet; initial development used small numbers of those studies (1-10 at a time) and we eventually developed a primary corpus of 400 studies that were confirmed to be curated to Gemma’s guidelines as completely as we could (50 of these had not been previously curated in Gemma). The 400 span mouse, human, and rat studies across a range of biomedical topics (**Supplementary Figure 3A and B**). The 400 are typical of Gemma, manually curated with 0-2 Experiment tags to fill in information not already captured by ontology-grounded sample characteristics, and 1-2 Factors (not counting technical batch factor), each with at least two Factor Values or numerical measurements (**Supplementary Figure 3C**).

The 400 can be considered the training data for this study, though as mentioned part of the Gemma corpus was also available to the system as reference data. We took steps to limit how much the pipeline could be contaminated by specifics of the 400 or be otherwise overfit to them. First, only parts of the data were examined at any iteration, at first only 1 or 5, and eventually 200 studies, as we iterated to improve the method. Only for the results presented here were all 400 processed by the v1.1 pipeline. Second, we actively prevented these studies being used as verbatim examples in the LLM prompts, instead using synthetic or generic examples.

The annotations of the 400 were frozen, as was the agent code base, and the pipeline run on them forming the basis of the “training” evaluation we report (due to unforeseen circumstances, the v1.0 pipeline had bugs that resulted in a major regression, which is why we report results for v1.1). We also report results for a test set of 100 test studies drawn from the 2,394 held-out, and thus unseen during the development process so far. While all 100 are in Gemma, they were only made public on our site after the training data cutoff for the most recent LLMs we tested (January 2026).

To simulate the task of a curator, we stripped the manually-added annotations from the Gemma data. We sometimes call this the *skeleton* or the *pre-curated* state of the experiment, as it represents what we receive when first loading the experiment into Gemma, including the automatically assigned ontology terms mentioned above. The task is to recover the stripped information.

### Proposer

The v1.1 proposer has eight main phases, seven of which involve one or more LLM calls (overview in **Figure 2A**). The prompt used for each phase was initially designed by the Claude coding agent, and iteratively improved by a range of approaches, including asking Claude to shorten prompts or emphasize certain aspects, or by having Claude generate multiple variations of the prompts to be evaluated. The proposer starts by making an overview of the study (“framing”), then deciding which features should be factors, identifying tags, and grounding everything in ontology terms (see next section). In some steps, purely deterministic analyses are used. In others, this can optionally escalate to an LLM inspection. The agent can use a “self-revision” component, which can lead to multiple cycles of internal review, optionally resulting in a re-call of the design or tag proposer with instructions on what needs further work. The final output is a JSON data structure that contains the proposed annotations and extensive meta-data from the pipeline: telemetry, justifications, and textual evidence including supporting quotes from papers.

**Figure 2:**
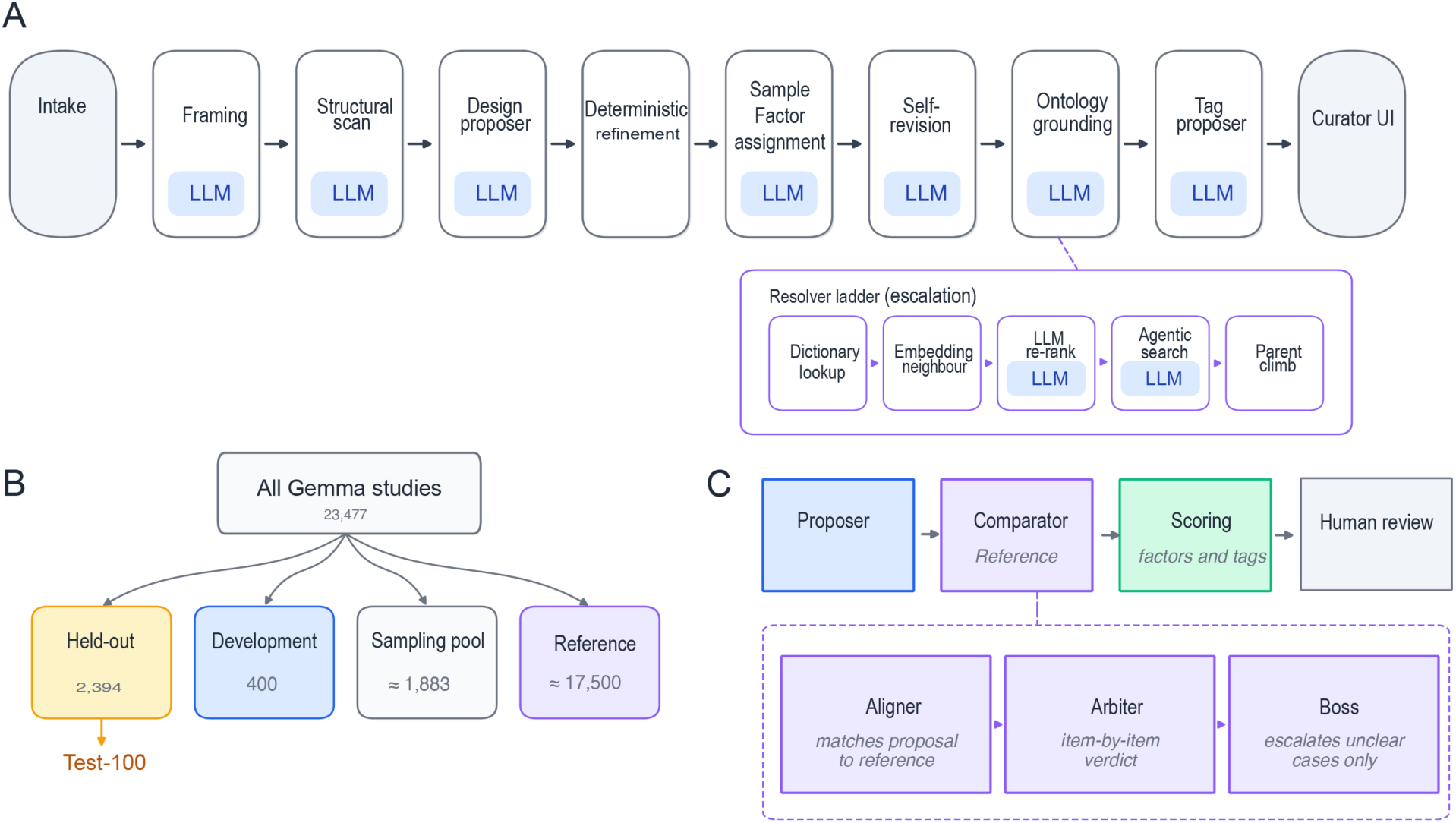
The proposal pipeline and evaluation framework. **A**: High-level schematic of the proposal pipeline. See main text and supplement for explanations. Intake of study meta-data is followed by an overall summary and framing, design (finding the factors, their values, and matching each value to an ontology term), whole-experiment tags, and a review pass, with the step-by-step term-matching cascade and an example of a matched factor value shown below it. **B**: Partitioning of the Gemma corpus as used in this study. Our primary analysis is on the 400 datasets, with a test on 100 studies held out during pipeline development. **C:** Post-proposal review and evaluation strategy: The pipeline proposes a curation without seeing the reference. The comparator finds correspondences between the proposal and reference (alignment), and which is then scored at varying stringency for correctness of: factors, categories, tags and ontology term matching, with emphasis on accounting for the semantic equivalence ("there’s more than one way to curate it"), evaluated by LLM calls. The "arbiter" and "boss" together play the role of an auditor; their feedback is provided to the manual reviewers in the Gemma curation interface (**Supplementary Figure 2**)

### Ontology grounding

We developed an ontology term and gene resolver chain that builds on our past work (Rogic et al. 2026), which is shared across the pipeline. This is backed by Gemma’s ontology search web services, which can use term frequency in Gemma annotations to help promote terms (for evaluation, the effect of the experiment being annotated was subtracted from this to avoid bias). This yields the highest performance to date on the benchmark of Rogic et al. (2026) with a ∼4-percentage point improvement for mouse strains (81% versus 77% correct) and cell lines (62.8% versus 59%). A further core design principle of the pipeline is that ontology terms (labels and URIs) emitted by LLMs are not trusted, so they all are subject to inspection and validation by mechanical and LLM stages. These steps are in addition to the resolver, and the other review phases can also result in corrections before final output.

### Post-proposal review

We anticipated that Gemma’s existing curation would not be a gold standard from the start. Not only have our guidelines changed over time and the data not perfectly kept up, human curators make errors. Most commonly these are omissions (e.g. failure to annotate that the animals were all male) but more rarely factual (e.g. mixing up two genes). Importantly, two “disagreeing” curations can both be correct, or at least acceptable. This creates a challenge for evaluation; the goal is to provide accurate, useful annotations, and our guidelines do not determine the outcome of every decision the curator has to make. Care was needed to distinguish errors from synonyms or semantically-close annotations which would otherwise be counted as both a false positive and a false negative. Similarly, proposals that appear to be false positives often turned out to be just gaps or over-annotation in Gemma; that is, the agent was correct.

The job of the post-proposal steps (**Figure 2C**) is to make identifying these nuances as easy as we could. The *aligner* component of the pipeline takes as input two sets of takes as input two sets of annotations, e.g. an agent’s proposal and gemma’s current curation, and identifies initial proposed correspondence or discrepancies between them, without using LLMs. The next phases play the role of a critic. This is distinct from the internal self-revision the proposer can do, and that the entire proposal is reviewed against criteria to check for quality and correctness, and (for this paper) this can be done in the context of comparing the existing curation (aligned). The input is the output of the aligner phase described earlier. The arbiter, makes an item-by-item judgement as to which of the two sets of annotations is correct, which (for the evaluations in this paper) we then treated as input to the humans evaluating the agents’ output. Cases the arbiter cannot confidently resolve are escalated to a second pass, which we call the “boss”, using a stronger model with extended thinking; by construction this is a minority of cases. See the interactive pipeline diagrams in the code repository for further detail on this and the other stages. The eventual intention is that this phase would be used to audit existing curation compared to the agent proposer, but we use it here for evaluation. We also plan to evaluate a more complete audit mode.

### Evaluation

Two human curators (authors CY and AM) actively participated (with PP serving as a third ad hoc curator and evaluator) in the pipeline development process. They used the web interface we developed to evaluate proposals made by the pipeline, and to accept or modify them, and along the way “polish” (i.e. bring up to current standards) the Gemma curation. After some small trials, there were several major rounds of human evaluation as part of development, first with 50, then 200 studies, and after some additional iterations to improve the system based on the feedback, the second set of 200 were then evaluated by both. Final decisions at points of disagreement were determined by re-review and final decision-making by PP where needed. This yielded the 400 “development” studies. The same process was carried on the test 100 data sets, ultimately including the polishing to bring them up to modern Gemma standards.

### LLM selection

We designed the framework to be able to use different LLMs at different stages, in an effort to save time and cost by not using “too-strong” models where they weren’t beneficial. We divide models into three tiers, “fast”, “standard” and “strong”. The latter two models must have the ability to provide structured output, and the strong model must have reasoning features. In v1.1, these are Haiku 4.5 (except in the tag proposer where it was Sonnet 5), Sonnet 5 and Opus 4.8. We report preliminary findings for a range of model choices for different stages of the pipeline, and evaluated an open weights model (GLM 5.2) (Team 2026). We also devised a version of the pipeline (“no-LLM”) that only uses the parts that do not use LLMs. This was not to test how well a non-LLM method could do, but to document the contribution of the LLM. There is no non-LLM route in our pipeline that proposes experiment-wide tags, so this baseline covers only Factors. This reflects how we built the mechanical steps, not a limit of non-LLM methods: the tags we score describe constant whole-experiment properties (disease, study design, disease model) that the deterministic proposer, working from sample-level metadata, does not attempt to infer. We leave the question over whether a non-LLM method could contribute more to tag generation.

### Publication identification subtask

Similar to the experimental design curation task, we provide the proposer with skeletons of experiments but this time removing the publication information from both the skeleton and the GEO record that the proposer retrieves. This simulates a common situation: the data set is public, it has been published, but the GEO record has not been updated to capture this information. The proposer uses a strategy that matches what curators do, which is to do some web detective work: Pick key words from the title and abstract and search PubMed; if that fails, search more broadly and even use the data submitter’s name and/or email address to find the laboratory they work in and search that person’s web site, where papers sometimes appear. We use biolit (github.com/rachadele/biolit/) to retrieve full text articles. Some remaining missing articles for the evaluation data were filled manually with the help of Zotero (zotero.org).

### Validation of outputs

A concern in relying on LLM-generated code is the possibility of errors or even so-called cheating. For example, we asked the coding agent to strip curator-generated information prior to putting the experiment through the proposer. If it had failed to do so, it could easily leak information into the process that would bias performance upwards. We addressed this sort of concern in several ways. First, prompts and code were repeatedly reviewed, both by coding tools and by humans. The agent code repository has over 3,400 tests, including end-to-end tests that verify the integrity of the pipeline. The evaluation process itself is implemented in python scripts. Second, inputs and outputs were scrutinized as one would for any data analysis. No cases of agent-caused information leakage were found, and in fact instead we identified (and addressed) agent errors or bugs that hurt performance (we take full responsibility for the code, data and results). Third, humans were heavily involved in doing performance evaluation by hand, through the UI, without noticing any anomalies. If the agent was smart enough to cheat, it would have been smart enough to do the curation more correctly. Finally, by making our code and data open anybody can audit our work.

### Triage analysis

We developed preliminary models to predict which agent outputs were likely to contain errors, to the end of prioritizing which would need human review. Errors of commission and omission were handled by separate models, both evaluated by dataset-grouped five-fold cross-validation. Full details are in the Supplementary Methods. Briefly, for factual (commission) errors we trained a gradient-boosted classifier (Pedregosa et al. 2011) on per-annotation features such as the consistency between a factor’s category and its ontology branch, resolver confidence and margin, and signals from the review chain, with confirmed errors as the positive class; it was evaluated by dataset-grouped five-fold cross-validation and we report the out-of-fold area under the ROC curve (AUC). For omissions, which have no emitted annotation to score, we worked at the level of the whole experiment, scoring each experiment by a severity-weighted mass of its differences from the reference. We related this score to an LLM-derived numeric representation of curation difficulty; associations are reported as Spearman correlations.

### Metrics

We report performance at several levels of granularity, as the curation task involves multiple decisions: whether a feature (e.g. an experimental factor) was detected at all, what category it belongs to, what ontology term its values map to, and what tags apply to the experiment. Our guidelines are flexible enough permit more than one valid way to express the same concept, so we report so results at more than one level of stringency.

1. Factor detection: For each dataset we computed recall as the number of reference factors also proposed by the agent out of total reference factors. A reference factor counted as detected if the agent proposed a factor with a matching category (for example, both labeled "genotype"), regardless of whether the factor’s values were later scored correctly.
2. Category assignment: A detected factor received full credit if its assigned category matched the reference category, and half credit if the two categories were both defensible for the same underlying concept (for example, "disease" vs. "disease model"). All other category assignments received no credit.
3. Factor value ontology mapping: We scored ontology proposals at three levels of stringency. *Exact* credited only an identical ontology term. *Graded* also credited a parent or child of the reference term, i.e. a coarser or finer term along the same ontology lineage. *Acceptable* further credited a sibling or cousin term close enough that a curator would likely consider it permissible. A mapping to an unrelated or absent term failed all three levels. We report precision, recall, and F1 at each level.
4. Whole-experiment tags: Tags were matched by exact (category, value) agreement between agent and reference and scored using F1 as above.

### Statistical analysis

Confidence intervals on the summary scores in Figure 3 are 95% Wilson intervals based on the number of items scored (experiments, factor values, ontology mappings or documented fields, as appropriate). We used replicate runs to estimate noise due to LLM variability and treated effects more than two standard deviations of this band as potentially significant. Curator agreement, shown as reference lines in Figure 3, is the agreement between two independent curators on the same studies. To test whether a dataset’s category predicts how often the agent’s factors differ from the reference we used ordinary least squares regression, and where we report an association between two per-dataset quantities we use the Spearman rank correlation. Analyses were carried out in Python.

**Figure 3:**
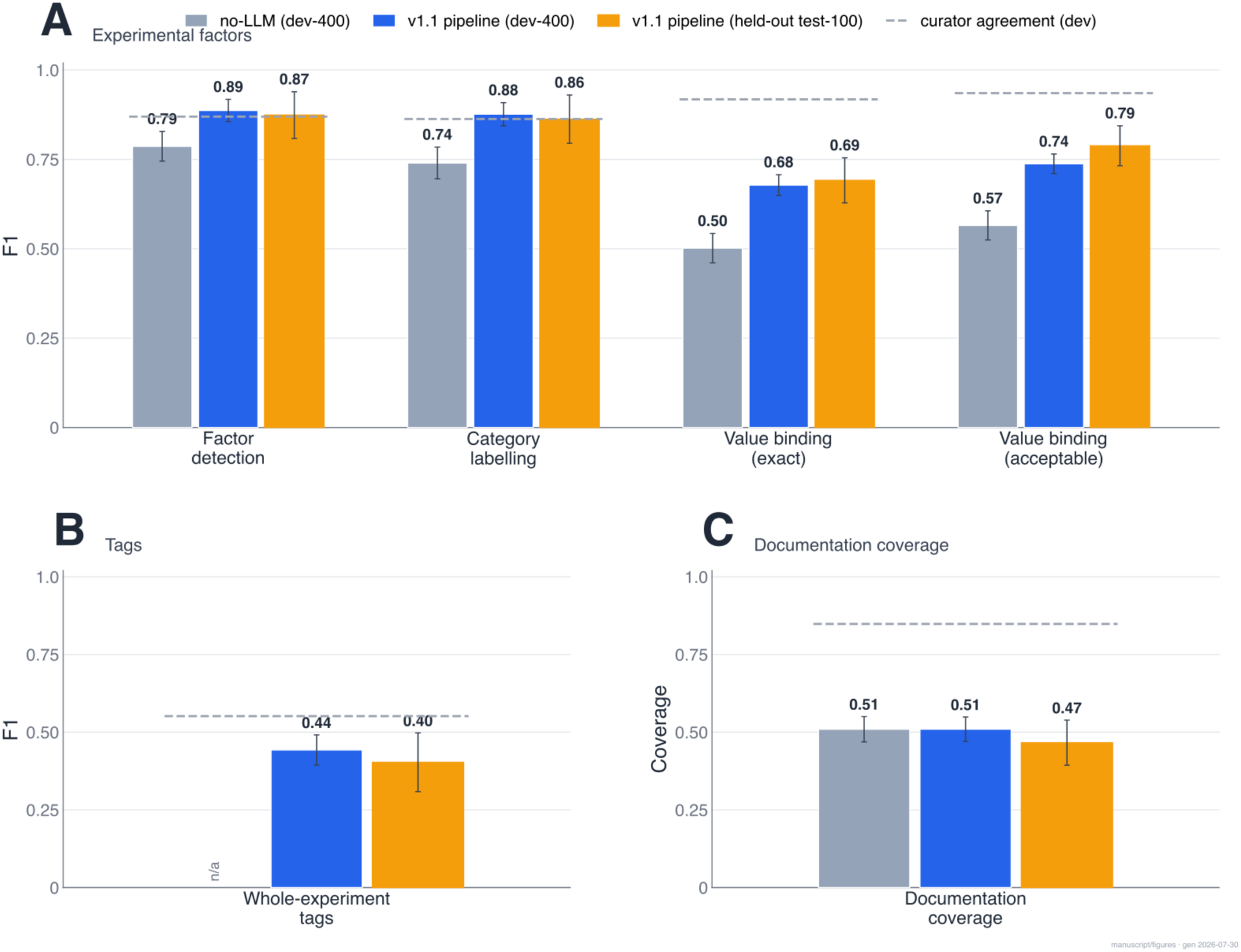
Primary performance results. The v1.1 pipeline is compared against a no-LLM baseline (grey), for the 400 and the test 100. The gap between the no-LLM baseline and the pipeline is the language model’s contribution; the closeness of the development and held-out bars shows whether performance carries over to unseen data. The dashed grey lines indicate the agreement between two human curators. Error bars are estimated 95% confidence intervals. **A**: Experimental factors, whether the right factors were found, whether each was placed in the right category, and whether its values were bound to the right ontology term at two levels of strictness (an exact match, and a more permissive level that also credits a closely related term). **B**: Whole-experiment tags. Note that the no-LLM baseline has no mechanism to propose these tags. **C**: Documentation coverage, the fraction of standard curation fields documented with an ontology-anchored value.

## Results

The task studied in this paper is curation of transcriptomics data at the experimental design and sample levels, intended to supplement or even replace the current manual process in Gemma. **Figure 1A** gives an overview of Gemma’s overall processing, of which the design curation task is one major step. **Figure 1B** is a schematic of the design curation task at a high level. The basic scheme is simple, but establishing these annotations is not easy. One challenge is that the GEO metadata is messy, and it requires integration of complex information and good knowledge of many areas of biomedical research to understand studies and convert the understanding into (more) formal annotations. Often critical information is in the text of a paper, or in a cryptic sample name. The second challenge is that our curation follows a complex scheme, with correspondingly complex rules and guidelines involving many large ontologies. Creativity can be needed to express a concept in the confines of our curation framework. Curators need months of experience to become proficient and much longer to be expert. It takes anywhere from 15 minutes to an hour to annotate a study without any special LLM assistance, though curators now use LLM chatbots or LLM-enhanced search (e.g. Google) in the course of their work. The curation task also varies widely in difficulty from data set to data set.

We developed a software pipeline, which we refer to as the **Gemma curation agent**, that mimics the job of a curator. **Figure 2A** is a schematic of the curation agent (version 1.1). A typical run for one data set will result in about 25 LLM calls, but there is an extensive software harness (what we refer to as the pipeline) which includes many “mechanical” steps that do not depend on LLM calls, or at least are only escalated to call an LLM when necessary. Each decision made by the pipeline comes with provenance information and justifications such as quotes from the paper or GEO meta-data. See the Methods and code repository for more details.

For evaluation, we ultimately developed a benchmark set of 400 studies (9,859 samples), picked randomly from ∼23,500 in Gemma, biased somewhat to more recent studies (**Figure 2B**; Gemma started curating data in 2005). Among these were 50 studies that had not yet been curated in Gemma, so they were curated during pipeline development. The annotations of these 400 data sets were thus reviewed and improved (“polished”) to bring them up to our current curation standards. Because these data sets were used to develop the pipeline, we refer to them as the development 400, but they can be thought of as training data. A further set of 100 previously unseen data sets were used for final testing (test 100). The properties of the data are summarized in **Supplementary Figure 3**.

**Figure 3** shows the primary findings for the 400 data “training” data sets and the 100 “test” data sets, for the two main tasks at a range of granularities and strictness. We plot the approximate level of curator performance (based on the human evaluation here) to help set expectations. We stress that it was a major evaluation challenge that two sets of annotations can be correct while differing (see Methods). The agent reproduced the reference curation *exactly* for only 30/400 datasets (7.5%), but far more would be judged “acceptable”. Our metrics attempt to reveal this. **Figure 4** shows a more detailed analysis of deviations between the agent and the reference annotations. The agent made many decisions that, when compared to the existing curation, led to the correction of errors in Gemma, filling in of gaps, and better standards adherence.

**Figure 4:**
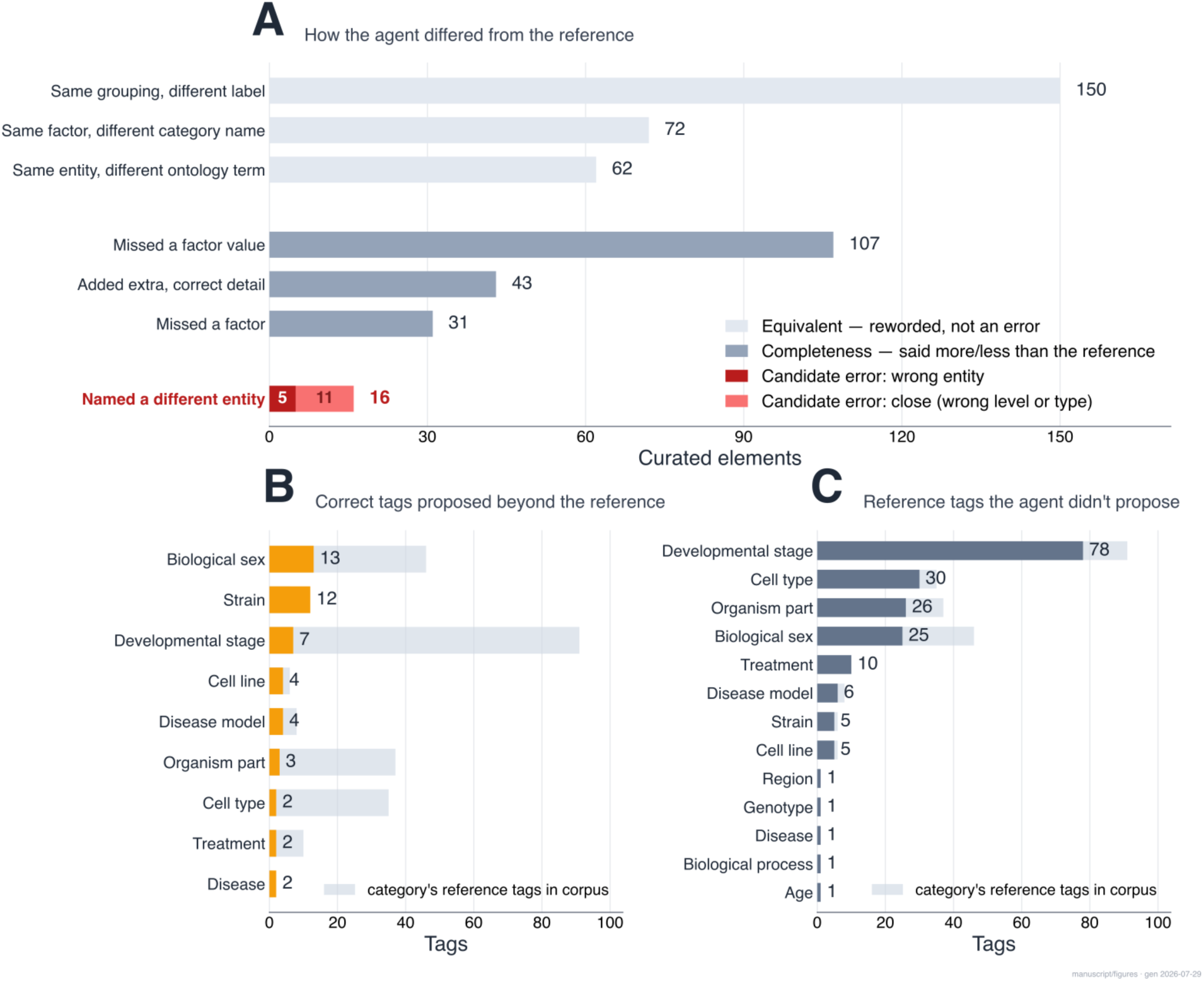
Differences between agent and Gemma curation. **A**: Overview of all types of difference, grouped by severity. The lowest severity (top, light grey) represents the same grouping, factor, or entity expressed under a different label or ontology term; we don’t consider this an error. The middle group (grey) is a difference in completeness: one side stated more or fewer details than the other. The bottom group (red), naming a different entity than the reference, are considered factual errors and of the greatest concern. **B**: Categories of correct experiment tags added by the agent but which were not already annotated in our original annotations (49 total). These were counted as true positives and the gold standard now reflects them. **C**: Tags the agent failed to propose, by category (false negatives) (190). In B and C, the grey bars indicate the number in each category in the corpus.

The performance is better than we expected to achieve at this stage of development, as the pipeline is far from fully optimized. A major theme of the findings is that serious errors were very rare, by which we mean “factually wrong in a way that would be a concern for downstream analysis or interpretation”. The pipeline almost always finds that an experimental factor exists (recall 0.98), but recovering the precise factor values is more challenging (F1 0.62), and whole-experiment tag performance is lower still (F1 0.45 against a curator ceiling of about 0.55). But crucially, the issue was often a failure to precisely follow our curation guidelines, rather than a factual error: the agent was trying to do the right thing but messed it up a little. In particular we see room to improve the ability of the agent to properly express concepts as statements. Recall of tags is also a gap, but omissions have lower impact, in our judgement. Results on the test 100 were very similar, suggesting that we have not grossly overfit to the 400.

Factual errors are the most concerning type of mistake so we took particular care to analyze them. On the 400, there were 16 potential such substitutions flagged, but on further inspection found only 8 to be genuine agent errors; the other 8 were cases where the agent was effectively right (a synonym or equivalent naming that our mechanical scorer did not yet recognize as a match) (**Figure 4A**). The most serious errors were confusing the gene being manipulated with the study’s background genetic model (GSE74438: a CD33 knockdown experiment in an APP/PS1 mouse background was annotated as an APP genotype) or naming the wrong kind of entity altogether (GSE31486: a CTCF knockdown was annotated with the cell line names BJAB/BL-41 instead of the gene). Conversely, the agent caught genuine errors in the existing Gemma curation itself, such as a Kit/Kitl mix-up (GSE277000) and a Rag1/Rag2 mix-up (GSE67136), which were corrected.

The pipeline is complicated, with at least 20 discrete components. To gain some insight into pipeline’s behaviour we plot a series of descriptive statistics in **Supplementary Figure 4**. One point of interest was that the internal critic (“arbiter”) mechanisms can result in multiple runs of some proposals, but our evaluation suggest that this was of only moderate impact. Indeed, of 860 arbiter verdicts scored on the 400-dataset benchmark, 50 (6%) were flagged as unclear and escalated to the boss reviewer, touching 44 of the 400 datasets (11%); most escalations (42 of 50) were resolved automatically (in a second round), leaving 8 that were unresolved and (in effect) left for review. The performance metrics and telemetry provide some insight into potential predictors of strong or poor performance on particular datasets, or to components that might not be pulling their weight. Overall pipeline efficiency, LLM cost, speed and openness are important considerations and the v1.1 pipeline is not fully optimized.

We conducted experiments to understand pipeline performance in more detail. As shown in **Figure 3**, a version of the pipeline that omits all LLM steps reveals the strong positive effect of the semi-structured meta-data provided in GEO records on the detection of experimental axes. Performance is higher at every step when an LLM call is included (**Figure 3**). We stress that the no-LLM method is just the pipeline with the LLM steps removed.

We next tested several variations of the pipeline on a random set of 100 studies drawn from the development set of 400 to inform model selection for each pipeline step. We first varied the type and strength of LLMs used within the model (e.g., using Opus throughout, or Sonnet as the “strong” model instead of Opus), and an open-weights model (GLM 5.2) (**Figure 5, top**). It turns out that the selection of models we used for v1.1 was high-performing but we are making some adjustments. For the latest (v1.2, unpublished) pipeline we have switched to Opus 5 as the strong model and Haiku for the tag proposer, without an obvious effect on performance so far, and we also had made a change from Sonnet 4.6 to 5 during development without observing much difference. Based on our preliminary and provisional test, GLM 5.2 was a strong performer on the factor task, but weaker on tags. If we take these numbers at face value, it’s possible that the pipeline can be further streamlined, but whether these differences persist with more analysis is a question we are investigating.

**Figure 5:**
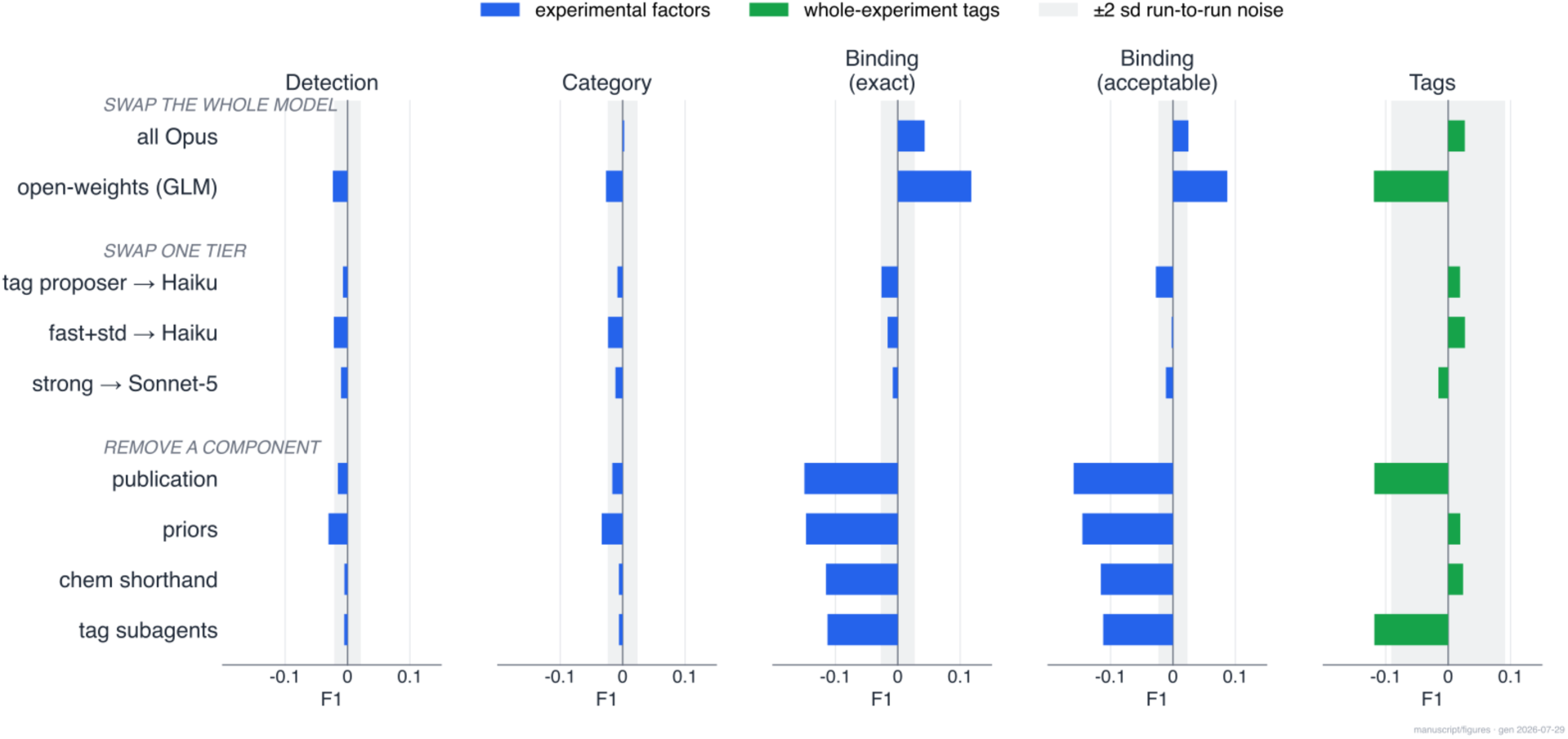
Contributions of pipeline components. Horizontal bars show how factor detection (blue; 261 factors, across four levels of granularity) and whole-experiment tags (green; N=54) change in F1 relative to the full pipeline, on 100 data sets randomly selected from the 400. The metrics are as in **Figure 3**. Each was run 1-3 times and noise estimates are from the pooled data. The *grey shading* shows the noise areas (2 standard deviations). **Top**, running the whole pipeline on a single model: all-Sonnet-5, all-Opus and an open-weights model (GLM-5.2). **Middle**: Effect of changing models for particular components. "Tag-proposer→haiku": The tag proposer has a LLM fallback that uses Sonnet-5 in v1.1. "fast+std→haiku": replace most Sonnet calls with Haiku; strong→Sonnet: replace most Opus calls with Sonnet. **Bottom**: The effect of disabling pipeline components. The components we have tested so far: *Chem shorthand*: removing the fallback that resolves chemical or drug abbreviations (for example DOX → doxorubicin) to ontology terms through a PubChem-to-ChEBI lookup, for shorthand the offline synonym tables miss; *Priors*: removing the knowledge drawn from previously curated Gemma datasets on common string-to-ontology-term mappings and how often each term has been used before; *Tag subagents*: removing the per-category tag specialists (biological sex, strain, cell line, developmental stage, organism part) that read the sample-metadata columns directly, and relying instead on a single monolithic tag-proposer call; *Publication*: removing the publication from the input. These are all non-LLM features except for some tag subagents that have LLM fallbacks.

We tested the impact the effect of other non-LLM components, such as removing the publication from the input (**Figure 5, bottom**). The presence of a publication was (on average) important to performance, especially for tags, agreeing with our experience. We hypothesize that occasionally supplementary files would also help, but we were not able to test this at scale. Other components we tested proved their value as well, such as the complex tag subagents outperforming the monolithic tag proposer. A component that proved to underperform was the self-revision within the proposer. The iterative loop for tags contributed almost nothing beyond its first pass: round 1 applied 248 of 260 tags (95.4 %), round 2 added 12 (7 correct, 3 false positives, 2 ambiguous) across 11 experiments, and round 3 added none, so although 233 of 400 experiments invoked a second or third round, the additional rounds were effectively net-neutral. The design self-revision layer was similarly inert. The latest (unpublished) version of the pipeline reduces the number of iterations. There are more variables that have not yet tested, but in general our ongoing attempts to improve the pipeline are focused on pipeline structure, prompts and the mechanical components.

### Prioritizing human review (triage)

If we could tell which of the agent’s curations most likely need fixing, we could forward only those and trust the rest. We developed simple classifiers (see Methods) to explore whether we can do this now. We have encouraging preliminary results for predicting errors of *commission*: a gold-blind classifier predicts whether a proposed annotation is a factual error with an area under the ROC curve of 0.86 (95% CI 0.74–0.90). (The test 100 had too few factual errors to evaluate). We note that LLM’s own stated confidence is actually anti-discriminative (dev-400 AUC 0.38), and a more useful signal is found in factors such as the consistency between a factor’s category and the ontology branch its value comes from (e.g., a tissue annotated under a genotype factor is a red flag). However, about 70% of the severity-weighted difference from the reference is errors of *omission*, and its largest component is low tag recall: the proposer under-extracts constant, paper-inferred properties such as developmental stage and biological sex. It is much harder to imagine being able to predict when something missing could be found. Still, we gave it a try. Omissions cannot be scored from emitted annotations, so we moved to the level of the whole experiment and looked for predictors of how much was missing; the best we found so far, an LLM’s own judgement of how hard the experiment was to curate, is only weakly associated with factor omission (Spearman ρ = 0.27; provisional test-100 results fared little better) and is redundant with raw scores derived from pipeline telemetry, so it is not helpful. We read this as a signal about where to spend further effort. We hypothesize that the most effective path is to make the proposer miss less in the first place, though triage remains worth developing alongside.

## Discussion

This work demonstrates that LLMs can be used to automate complex data curation tasks. While our work is ongoing at this writing, the number and severity of errors are already at, or near, what we feel is tolerable, and in some ways exceeds human performance. We are planning to implement further improvements and move the system into production for Gemma soon. While our system is site-specific and occupies a small niche of the biomedical experiment landscape, we learned many lessons that are likely to generalize to other tasks.

We want to be clear that we do not claim our method is high performing in some general sense: our comparisons are against our own human curators, not against other published systems, since no shared benchmark exists across these different tasks and datasets. Part of the challenge is that the field has not agreed on what counts as correct curation in the first place: what to annotate, in how much detail, and in what way, including which ontologies to use. At worst, though, we have provided a task that is a microcosm of that larger problem. We also do not claim that our approach of using agentic LLM for curation is novel in 2026, but we have not seen any other system that addresses the degree of detailed curation that we do here, whether LLM-based or not. The most elaborate of those mentioned in the introduction is probably the (closed-source) work of (Mondal et al. 2025), which annotates descriptive single-cell sample fields rather than reconstructing the experimental design. We are making our system and benchmark data available in the hope that others can benefit or in turn develop better methods. In the rest of the discussion, we comment on some of the lessons we learned, and our outlook.

Working with a coding agent allowed rapid iteration and the development of a system of a complexity that we would not have attempted otherwise, and we see this as one of the most important outcomes. It’s not that “old” methods cannot accomplish curation tasks to some degree, it’s that they were too costly to develop without knowing how well they are going to work ahead of time. The coding agent largely removes this barrier, allowing us to do ambitious and risky experiments with little cost, testing dozens of ideas in a matter of weeks. This “human in the loop” process was still labour intensive, and one might ask if a sufficiently capable agentic LLM could construct such a system by itself without human involvement, given only a specification. We did not attempt this (yet) and instead worked on relatively small pieces at any given time. The coding agent makes errors and forgets commonly used facts despite reminders. For example, we had many frustrating moments with Claude where it got confused about the relation between Experiment Tags and Biomaterial Characteristics, despite many attempts. We got better at durable remediation of such confusions, but it would not have been reasonable for us to pre-emptively write a specification that could have anticipated every possible snag. At the same time, that particular confusion was really of our own making: a long-ago decision about our data model that is no longer very relevant. It was an interesting exercise to try to get the coding agent and the curation to adhere to our curation guidelines, but an easier (if slightly uncomfortable) path is to adapt the task to the capabilities of the coding agent and underlying LLMs. For example, the agent tended to add annotations that we didn’t traditionally annotate, like the sex or age of the animals used in a study when this was a constant. This is consistent with our guidelines, but was never prioritized compared to features that vary. The agent gives us a benefit here, though as our results show it does not do this perfectly.

Besides writing code, LLMs contribute directly to agentic curation in two distinct ways. First, as a flexible text processor: agents handled complicated parsing challenges and spotted patterns in inconsistent metadata that would otherwise require complex text parsers and regular expressions for each case. Second, as a substitute for human judgement, synthesizing information from the curation guidelines and the disparate pieces of metadata associated with a study. As noted in the introduction, our goal was a system that could obviate most human curation, and our experience with both text processing and reasoning was generally encouraging. We experimented with multi-agent "back and forth" adjudication, and parts of the pipeline still work this way, but it was only partly effective. In general, adding another adjudication layer was usually less productive than improving the proposer directly, specifically by filling gaps in the information it had access to, improving prompts, and refining mechanical steps. It is notable that using models that rank lower on published general benchmarks (e.g., coding performance) had modest effects on the results and were notably not necessarily worse (**Figure 5**), which suggests LLM capability was not where we can improve right now, but larger evaluations may be necessary to make strong conclusions. We do believe that LLM performance is a limiting factor and developments over the next year or two could shift the picture.

The experimental annotation task, or at least our version of it, is difficult because it is complex. If nothing else, our attempts at automation show that asking an LLM to represent biological samples in an ontologically structured manner is non-trivial. It is difficult to even firmly define what is good annotation, in part because the types of experiments are almost endless. The flip side of this is our observation that while inter-annotator agreement is strictly limited (whether the annotator is human or a computer program), much of the time the difference is either one of “there’s more than one way to do it”, or arguably not very important. The stakes for our use case (perhaps not others) are not that high even for a factual error like mixing up two similarly named genes; such errors occurred with human curation too. We take the view that achieving perfect annotation is not possible much less practical, and that errors are going to occur. We must decide what detail and type of annotation we need, and types and severity of errors we can tolerate to accomplish what we want to do with the data.

### Future prospects

Financial support for data curation activities is difficult to obtain, and there is a large backlog. Is it worthwhile to clean up the old data? We are clearly biased in favour of one answer, but it depends on the value of the information, which for Gemma is the value of re-using the data. On average, most of the value of a GEO series is likely already spent on the analyses originally intended by the data producer and accessed via the associated publication. Any residual value is difficult to measure. It is dubious to put dollar values on data, but for the sake of illustration, if a typical genomics study of 10 samples costs $30,000US to generate (including reagents, samples, personnel, infrastructure and depreciation of equipment and facilities), a 1% ($300) residual value would be well worth it, even if curation cost $10 per study (our past approximate cost for Gemma). But if the residual value is nearer 0.1% on average, manual curation is harder to justify, especially at the scale needed. Computational annotation changes the economics dramatically if the error rates are tolerable, and we don’t want to be overly-optimistic about the errors, because it is still early. We also bear in mind that a major future consumer of curated data is going to be LLMs or other software systems, not individuals; this may also change the economics and considerations of the costs of errors. In any case, the v1.1 pipeline produces annotations at ∼$0.5 per data set. For Gemma, this moves the bottleneck from curation to computation (raw data re-processing; Figure 1A) and quality control, which is already heavily automated and has opportunities to be further encompassed by the agent. If 10% of studies still have to be examined by a curator we could move from a rate of 50 studies a week per curator (fairly typical, with ∼50% effort) to 500. Optimistically, this suggests we could re-curate all of Gemma in a year, or two years at an even lighter workload.

Besides re-curating old data, it is reasonable to focus efforts on data that is yet to be generated, by establishing frameworks for getting the curation right up front, rather than as an after-market add-on. Data repositories like GEO could require submitters to review an agent’s proposal for their study, at the time of submission. Schemes that have submitters do detailed technical curation have previously faced challenges, because a data curator has specific training and knowledge. Increasing burden on submitters reduces the chance they will submit to the resource in the first place. Human resources have to be spent to help submitters or else do it in their stead, which doesn’t scale. It is worth exploring whether LLMs can finally crack this problem, but we note that (as far as we know) GEO doesn’t do this type of detailed curation at all, leaving the problem more open-ended than for Gemma.

Another approach is encouraging the use of a curation agent in laboratories, as the experiment is being conducted or soon after rather than as an afterthought. If coding (or similar) agents become constant work-partners of researchers (as is the case in our laboratory), this could emerge very naturally and with minimal extra effort. This is because working with a coding agent tends to result in better documentation than a human working alone, at least relative to the effort from the human. A tool styled on our agent would encourage good data curation practices by implementing them just as readily as the agents write memory files or python scripts. Tasks like forming valid statements and finding ontology terms would be handled by the agent as a matter of routine. Unless a widely agreed annotation schema is developed this might not result in the level of uniformity we seek in Gemma, but it could be much better than nothing.

### Limitations

The Gemma curation agent is still a work in progress and we believe there are many ways to try to improve the agent and reduce errors; and at the same time, we expect more problems to crop up as we continue to test and apply it to more data sets. We know from experience that even after having seen 25,000 of them, every new GEO study has a chance of being a novel edge case. This is why effective triage will be important, so that only trouble spots need human review, which also feeds back to improve the pipeline.

While the experience and approach we developed may translate to other data types and settings, we have only worked with transcriptomic studies from mammals. But we are optimistic about generality because tissue sources, genotypes and so forth have little relationship to the assay that was run. Expansion to other species or different types of samples might require specialization, but hopefully not a complete rethink. We also do not claim that Gemma’s type and detail of curation is always what is needed, and our annotation model has limited expressivity. Those limits originated as compromises made for manual curation, but we suspect that if much more detailed or semantically-precise curation is desired, new difficulties will arise. Better support for complex genotypes is one specific gap we have not yet closed and now see the opportunity to tackle. With 500 studies that we have established as a benchmark, we have a means to test the effect of new improvements or to detect regressions. We note our evaluation was LLM-assisted, which despite our extensive manual inspection, could still introduce some errors or bias. But anecdotally, we found the LLM to be very helpful when dissecting both human and agent mistakes or near-misses.

In conclusion, we expect that systems like ours will soon advance to the point that tedious human curation of immense biomedical data corpora is neither economically nor humanistically justified. We are moving our pipeline for Gemma into production. We envision that with further work to expand its applicability, the curation agent could usefully curate all of GEO, even at current levels of imperfect performance.

## Contributions

PP: Project conception and oversight, development of the agent and user interface, contributed to data curation and evaluation; BOM: Contributed to software development; RAS: Developed the literature retrieval biolit MCP and contributed to manuscript development and editing. CY and AM: Curated data, evaluated the agent and contributed feedback on its development and that of the curation user interface.

## Acknowledgements

We acknowledge grants from the Canadian Institutes of Health Research (CIHR PJT-205846) and the Natural Sciences and Engineering Research Council of Canada (NSERC grant RGPIN-2016-0599) to PP. Additional computational resources were provided by the Anthropic AI for Science Program. None of these organizations had input to or influence over the study. We thank Dmitry Vavilov for system support.

## Conflicts of interest

The authors declare no conflicts of interest.

## LLM use statement

The research relied heavily on LLMs (see Methods). An LLM was also used to assist in manuscript preparation (figures, citations, formatting), and to provide initial drafts of some methods and results sections which were then heavily edited by the authors. Supplementary methods files and other documentation in the code repositories were largely LLM-generated. The ground source for the function of the system and the analyses presented is the code provided.

## Supporting data and software access

Gemma curation agent: https://github.com/PavlidisLab/gemma-curation-agents-v1.1; Gemma UI: https://github.com/PavlidisLab/gemma-ui. We note that to run the software fully requires access to a Gemma 2.0 REST endpoint, which is not public as of August 2026. **Supplementary Methods** are in the agent code repository. Benchmark data and v1.1 outputs: https://github.com/PavlidisLab/gemma-curation-benchmark-data. All software is open source (Apache 2.0 License). The benchmark data and Gemma’s curation are released under a CC-BY-NC license.

## Supplementary Figures

**Supplementary Figure 1:**
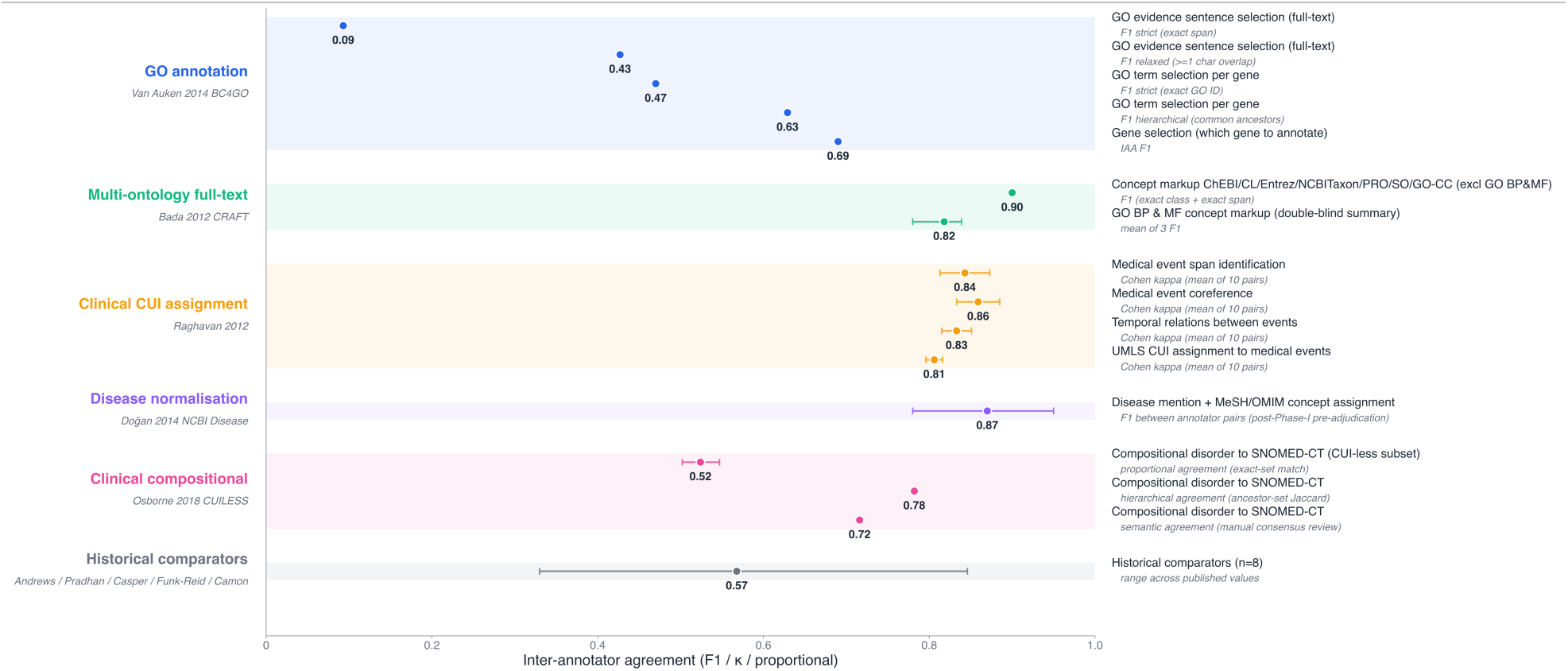
Inter-annotator agreement in biomedical curation tasks. A forest plot compiling published inter-annotator agreement across a range of biomedical curation tasks, from Gene Ontology annotation and disease normalization to clinical concept assignment. Each row reports the agreement metric that study used (F1, Cohen’s κ, or proportional agreement), so values are indicative rather than strictly comparable. Error bars are ranges or 95% confidence intervals, depending on the source. See supplementary references for full citations.

**Supplementary Figure 2:**
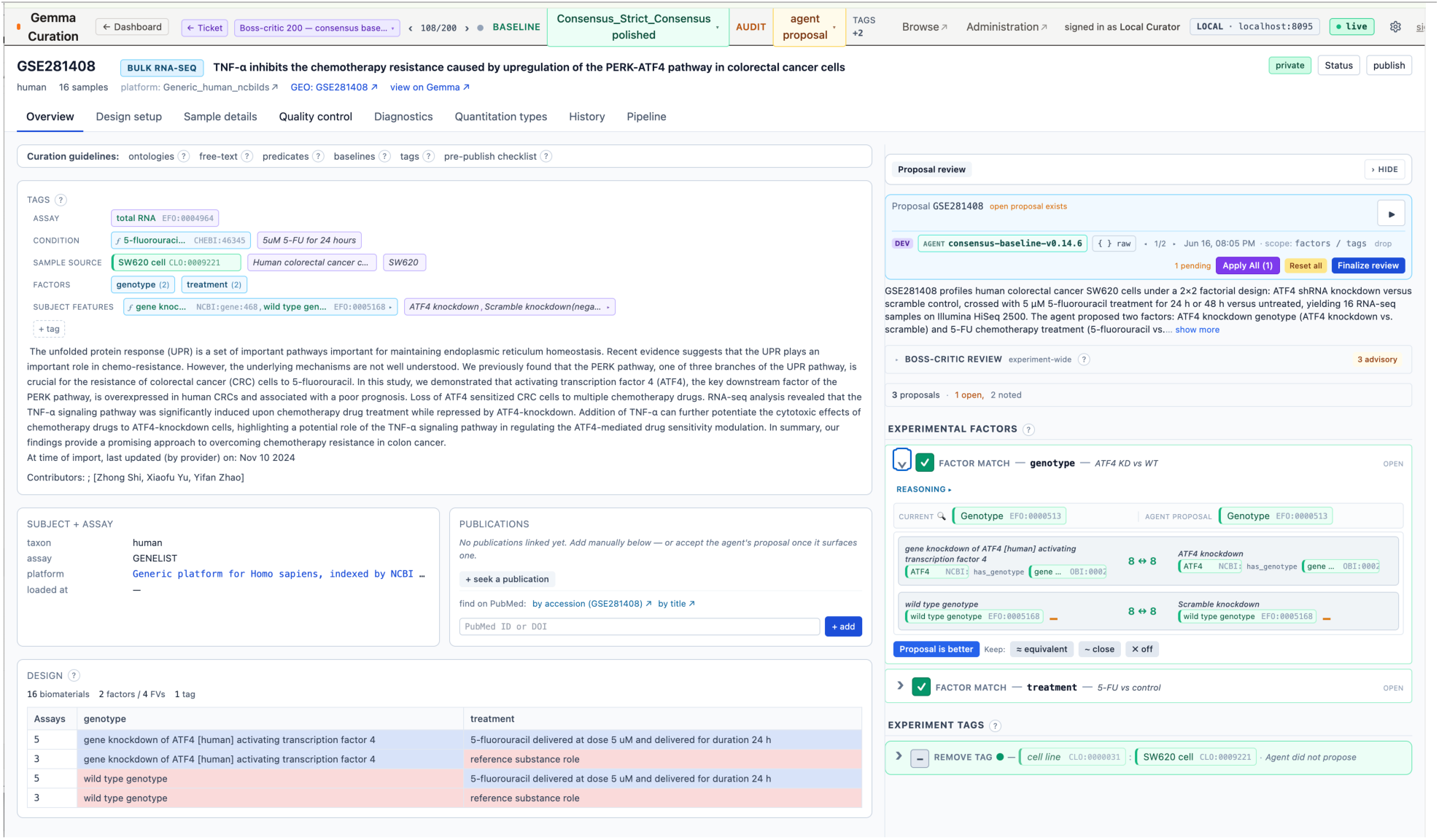
The curation interface with agent proposals side-by-side. A screenshot of the Gemma curation interface (example dataset GSE281408). The curator’s view of the study, left, is shown alongside a proposal-review panel, right, in which the agent’s proposed factors, factor values, ontology bindings, and whole-experiment tags appear next to the current curation along with the agent’s justification and evidence. Curators review computational proposals, making keep or reject decisions which are recorded by the system

**Supplementary Figure 3:**
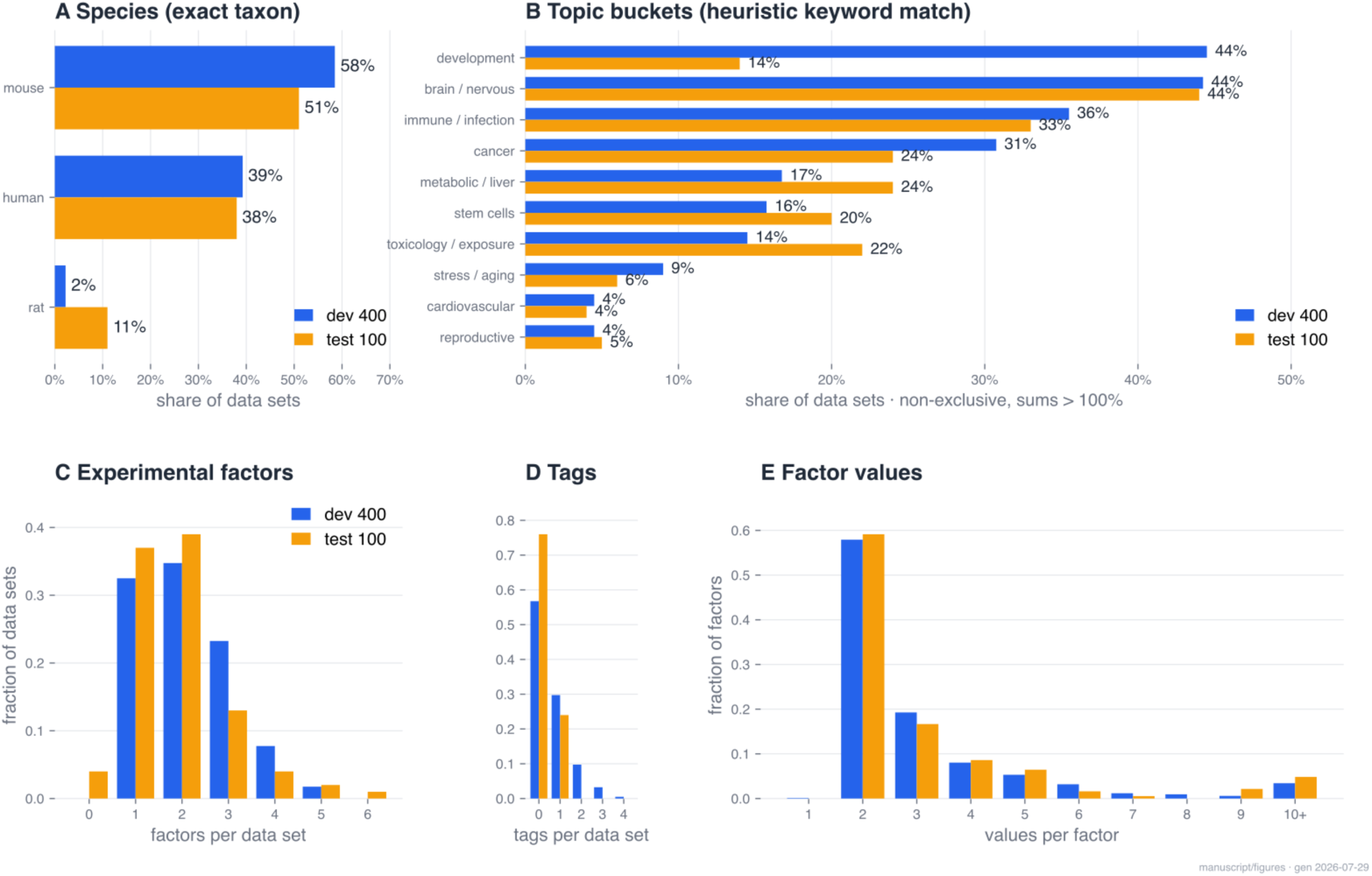
Properties of the benchmark and its reference curation. **A**: Species composition. **B**: Coarse biomedical topics, from non-exclusive keyword buckets, so a study can fall in more than one and the shares sum past 100%. **C**: Experimental factors per data set. **D**: Whole-experiment tags per data set. **E**: Factor values per factor, with the long tail binned. Every panel places the development 400 set beside the held-out test 100 set as a fraction of each, so the two compare despite their different sizes, though the test 100 happened to have fewer "development" studies.

**Supplementary Figure 4:**
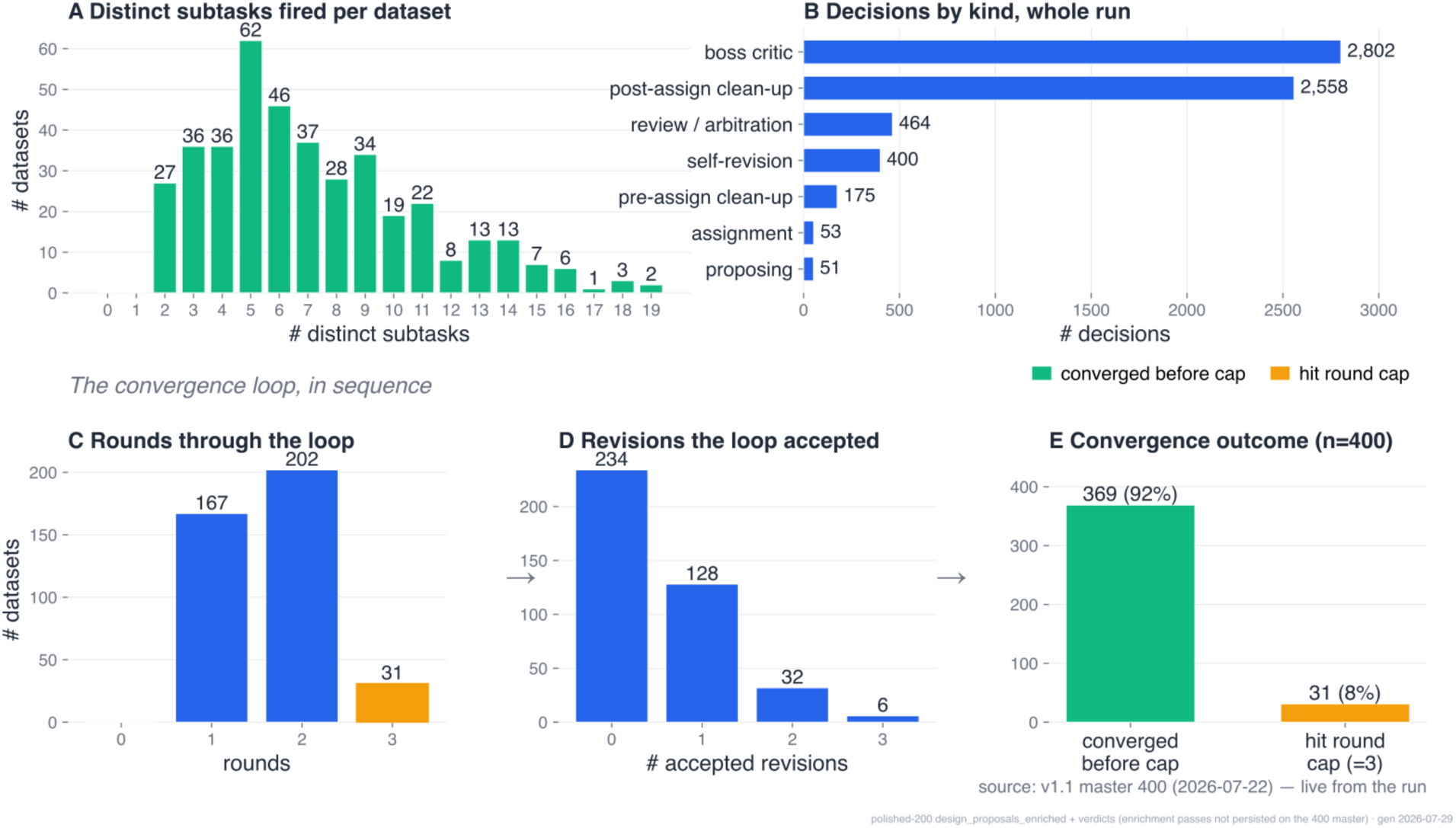
Pipeline behaviour across the 400 datasets. Because the pipeline chooses its own path, its behaviour varies from dataset to dataset. **A**: the number of distinct subtasks that fired per dataset. **B**: how many decisions of each kind the pipeline made across the whole run. **C–E**: Behaviour of the self-revision loop: **C** How many rounds each dataset took (the loop is capped at three rounds, the amber bar); **D** How many self-revisions the loop actually accepted; **E:** Whether the dataset converged before the cap or hit it.

## Notes

https://github.com/PavlidisLab/gemma-curation-agents-v1.1

https://github.com/PavlidisLab/gemma-ui

https://github.com/PavlidisLab/gemma-curation-benchmark-data

### Competing Interest Statement

The authors have declared no competing interest.

